# Phenomic selection enables nursery screening for field performance in maize and pea

**DOI:** 10.64898/2026.10.16.757273

**Authors:** R. Rincent, Q. Hamon, A. Garcia Dias, M. Ecarnot, L. Nunes, C. Bauland, P. Dai, A. Comar, G. Touzy, F. Barthes, D. Gouache, A. Le Mener, E. Goudemand, L. Moreau

**Author notes:** Corresponding author: Renaud Rincent.

## Abstract

Due to economical constraints, breeding programs eliminate most selection candidates at early stages, before their productivity can be evaluated in field trials. Genomic selection partially overcomes this limitation by enabling early prediction of field performance, but its implementation requires large-scale recurrent genotyping that remains inaccessible for many species and breeding programs. Phenomic selection offers a promising alternative by replacing genotyping with inexpensive high-dimensional phenotyping data. However, previous applications have focused on predicting genotypes already evaluated in field trials. We investigated whether phenomic data acquired by unoccupied aerial vehicles (UAVs) from nursery-grown genotypes can be used to predict their performance in field trials. We validated this nursery-screening approach in maize and pea, two species with contrasting biology and canopy structures, supporting its potential transferability to other species. In the most challenging scenarios, average accuracies for yield-related traits were between 0.36 for maize and 0.40 for pea. Simulations indicated that these accuracies can support substantial nursery culling while limiting the risk of discarding superior genotypes. This proof of concept demonstrates that UAV-based phenomic selection can predict the field performance of nursery-grown genotypes without genotyping the selection candidates.

## Introduction

At the early stages of breeding programs, thousands of newly generated genotypes are subjected to strong selection despite limited information on their agronomic value. Selection in nurseries is generally based on traits such as plant height, flowering time or disease symptoms, whereas productivity, usually the main breeding target, cannot yet be evaluated reliably. Because genetic gain depends on both selection intensity and selection accuracy^1^, applying strong selection pressure with low accuracy limits genetic progress and creates a substantial risk of discarding superior genotypes.

Genomic selection partly overcomes this limitation by enabling field performance to be predicted before candidates are evaluated in multi-environment trials^2^. Genomic predictions can be used to decide which candidates progress to field evaluation or, when accuracy is sufficient, to select parents directly for the next breeding cycle^3^. However, routine genomic selection requires recurrent genotyping of thousands of candidates every year. The associated costs and infrastructure requirements limit its implementation in minor and orphan crops^4,5^, in resource-limited breeding programs, and in regions with limited access to genotyping facilities^6,7^. More than 500 plant species are bred worldwide^8^, yet only a few dozen have a dedicated single-nucleotide polymorphism (SNP) array^9^, the standard tool for genomic selection. Genotyping-by-sequencing offers an alternative^10^, but it has limitations related to reproducibility^11^, imputation^12^ and genotype calling^13^, and is not always cheaper^14^. This gap matters beyond individual programs: the lack of improved varieties for most species is a recognized obstacle to crop diversification^15,16^. Predictive approaches that do not require genotyping every selection candidate could therefore complement genomic selection and broaden access to predictive breeding.

Phenomic selection provides such an alternative by replacing molecular markers with high-dimensional phenotypes (HDPs), such as near-infrared (NIR) spectra^17^. Because HDPs are partly under genetic control, they can potentially capture genetic similarities among individuals. They can therefore provide predictive information for complex traits, including traits unrelated to the tissue from which the HDPs were measured^17,18^. For instance, NIR spectra acquired at a single location predicted grain yield in an independent multi-environment trial as accurately as genomic prediction^17^. Phenomic selection has since been evaluated in cereals^19–28^, oilseed and protein crops^29–31^, forage crops^32,33^, fruits and vegetables^4,34–36^, and forest and fruit trees^17,37^. In these studies, accuracies were often comparable to those of genomic selection.

However, previous applications have largely focused on material already evaluated in field trials. To our knowledge, phenomic selection has not been evaluated for screening candidates while they are still in the nursery, before any productivity data exist, although this is arguably where it could have the greatest impact. The structure of breeding programs makes such screening feasible. In a given year, the genotypes evaluated in field trials are generally also grown in the nursery for seed multiplication. HDPs can therefore be acquired under the same nursery conditions for two sets of genotypes: those that are also evaluated in the trials, which provide a training population with field agronomic data, and the new selection candidates grown exclusively in the nursery. The phenomic similarity between the two sets could then be used to predict the field performance of the new candidates without genotyping them (Fig. 1).

**Figure 1:**
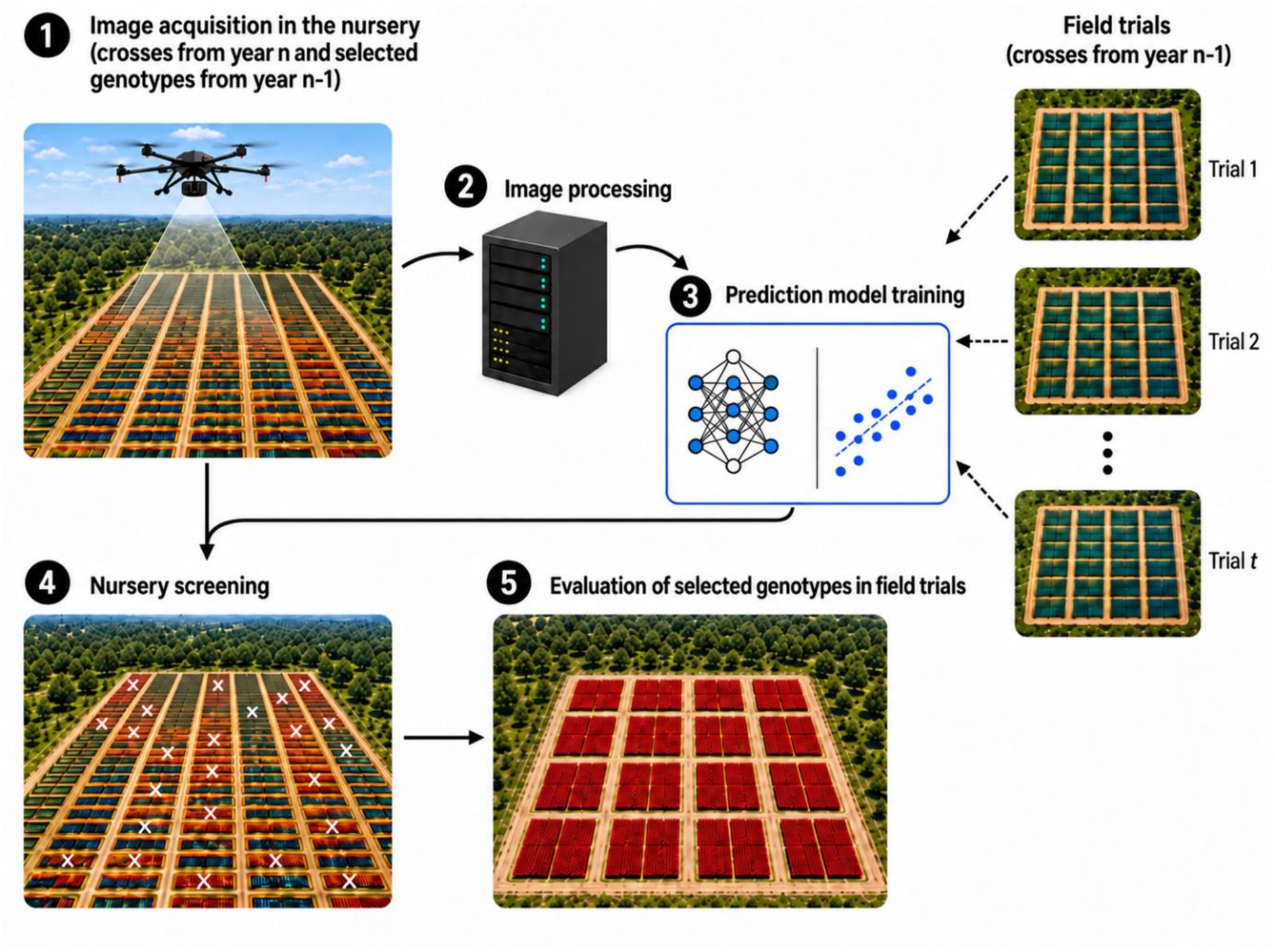
Conceptual framework of phenomic selection for nursery screening. For a given year, high-throughput phenotyping data is acquired in the nursery (1) and pre-processed (2). The resulting variables are used as predictors (as such or to estimate a similarity matrix between genotypes) in a phenomic prediction model (3). The training of this model is possible because all genotypes evaluated in field trials that same year are also present in the nursery for seed multiplication. The predictions are then used to screen the nursery and select the best genotypes for evaluation in field trials the next year. For hybrid crops such as maize, nursery material consists of parental inbred lines, whereas field trials evaluate hybrids. In this case, phenomic data acquired from the parental lines can be used to predict the performance of their hybrids, as demonstrated in this study.

Applying this approach at nursery scale requires phenomic methods compatible with thousands of plots. NIR spectroscopy has provided effective predictors for phenomic selection ^17^, but plot-by-plot acquisition becomes limiting in large nurseries, especially for species like maize, for which NIR is not used in routine at this stage. Unoccupied aerial vehicles (UAVs), in contrast, can image entire nurseries repeatedly throughout plant development. UAV imagery can be summarized as secondary traits describing canopy development, phenology, architecture, reflectance and biomass. It can also be represented by latent features extracted with self-supervised models pretrained on generic video, which require no annotation^38^. UAV-based phenotyping has been widely used to predict agronomic traits of the plots imaged in field trials^39–41^. Whether imagery acquired in nurseries can predict performance in independent field trials remains unknown. The nursery-screening approach based on phenomic selection that we propose here is challenging for two reasons compared with previous work, including our recent study on maize^26^: first, growing conditions differ substantially between field trials and nurseries in experimental design, sowing density and management, with genotypes grown in rows in nurseries and in plots in trials; second, to our knowledge, UAV imagery has not previously been used to predict field-trial performance from nursery-grown plants.

Here, we tested whether phenomic data acquired in breeding nurseries can predict the field performance of unevaluated selection candidates. In maize, we evaluated whether NIR spectra and UAV imagery acquired from parental lines grown in nurseries could predict the productivity, phenology and quality of the resulting hybrids in multi-environment trials, and we compared these predictions with genomic prediction. We assessed prediction accuracy when the parental families of predicted hybrids were absent from the training population, and we examined the effects of UAV acquisition timing and number of flights. We then tested the approach in a winter pea breeding nursery. Pea differs from maize in its biology (dicotyledonous and self-pollinating versus monocotyledonous and cross-pollinating), breeding system (inbred lines versus hybrids), crop status (minor versus major crop) and canopy architecture (low, dense and spreading versus tall and upright). Finally, we quantified how prediction accuracy affects the risk of eliminating superior candidates during nursery screening.

## Results

### Study design

We tested nursery-based phenomic prediction in two species (Fig. 1). In maize, the dent and flint parental lines of an incomplete factorial design were grown in two nurseries in 2024. Near-infrared (NIR) spectra were acquired from leaves and grains, and UAV imagery was acquired on six dates spanning the season, from emergence to senescence. The UAV images were represented in two ways: 198 secondary traits describing canopy development, reflectance and biomass, and 1,280 latent variables extracted with a self-supervised video model (V-JEPA) used without fine-tuning.

These nursery data were used to predict the performance of 242 dent × flint maize hybrids, derived from 68 dent and 77 flint parental lines. The hybrids had been evaluated in eight field trials across multiple environments in 2016 and 2017. Because the nursery was grown several years after the hybrid trials, predictors and target phenotypes share no environmental effects. Target traits were dry matter yield (DMY), dry matter content (DMC), silking date and milk fodder unit (MFU), and phenomic predictions were compared with genomic prediction based on 9,548 SNPs^42^.

In winter pea, UAV imagery was acquired on four dates in a breeding nursery of 464 genotypes from 100 F4 families. The nursery imagery was used to predict yield, plant height and flowering date of the same genotypes in a field trial conducted the same year. Throughout, prediction accuracy is the Pearson correlation between predicted values and adjusted phenotypic means of field performance, divided by the square root of the trait heritability (see trait heritabilities in Supplementary Note 1).

#### Nursery UAV imagery of parental lines predicts hybrid performance

We first used a random cross-validation scenario (CV_related: 80% training, 20% validation, 200 replicates). In this scenario, training and predicted hybrids can share parents and parental families. It therefore provides an upper benchmark, more favourable than most breeding situations.

Genomic prediction reached a mean accuracy of 0.88 across the four traits (Fig. 2). UAV-based phenomic prediction came close, with a mean accuracy of 0.82 (UAV secondary traits), whereas predictions based on NIR spectra were less accurate. Combining the two NIR sources (leaf and grain) or the two UAV representations (secondary traits and latent variables) increased accuracy in most cases. This indicates that the predictor types carry complementary information.

**Figure 2:**
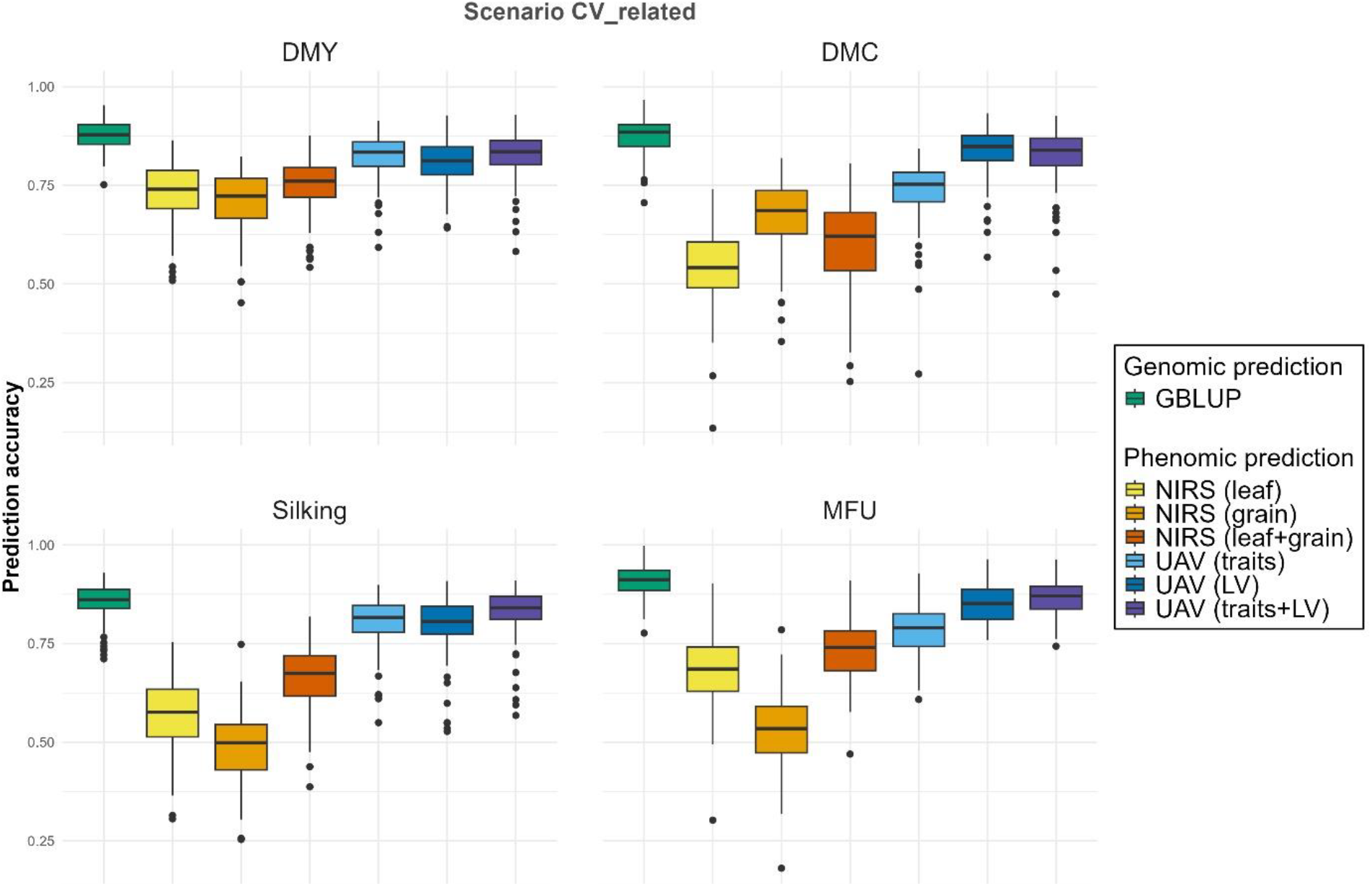
Accuracy of genomic and phenomic predictions obtained in random cross-validations among the 242 maize hybrids (scenario CV_related). In this scenario, the training and predicted genotypes are strongly related. Predictions of hybrid performance were based on near-infrared spectroscopy (NIRS) or UAV images acquired in the nurseries on the parental lines. DMY: dry matter yield, DMC: dry matter content, MFU: milk fodder unit.

We next considered a more stringent scenario (CV_newFamilies; 150 partitions). Here, the parents of predicted hybrids belonged to families absent from the training population (Fig. 3). Accuracy decreased substantially relative to CV_related. Mean accuracy across traits was 0.41 for genomic prediction and 0.33 for phenomic prediction based on UAV secondary traits. For UAV secondary traits, accuracy ranged from 0.20 for DMC to 0.48 for MFU, and reached 0.36 for DMY. For DMY, the difference between genomic and UAV-based prediction was 0.07. Latent variables were less predictive than secondary traits in this scenario, and combining the two UAV representations did not improve accuracy.

**Figure 3:**
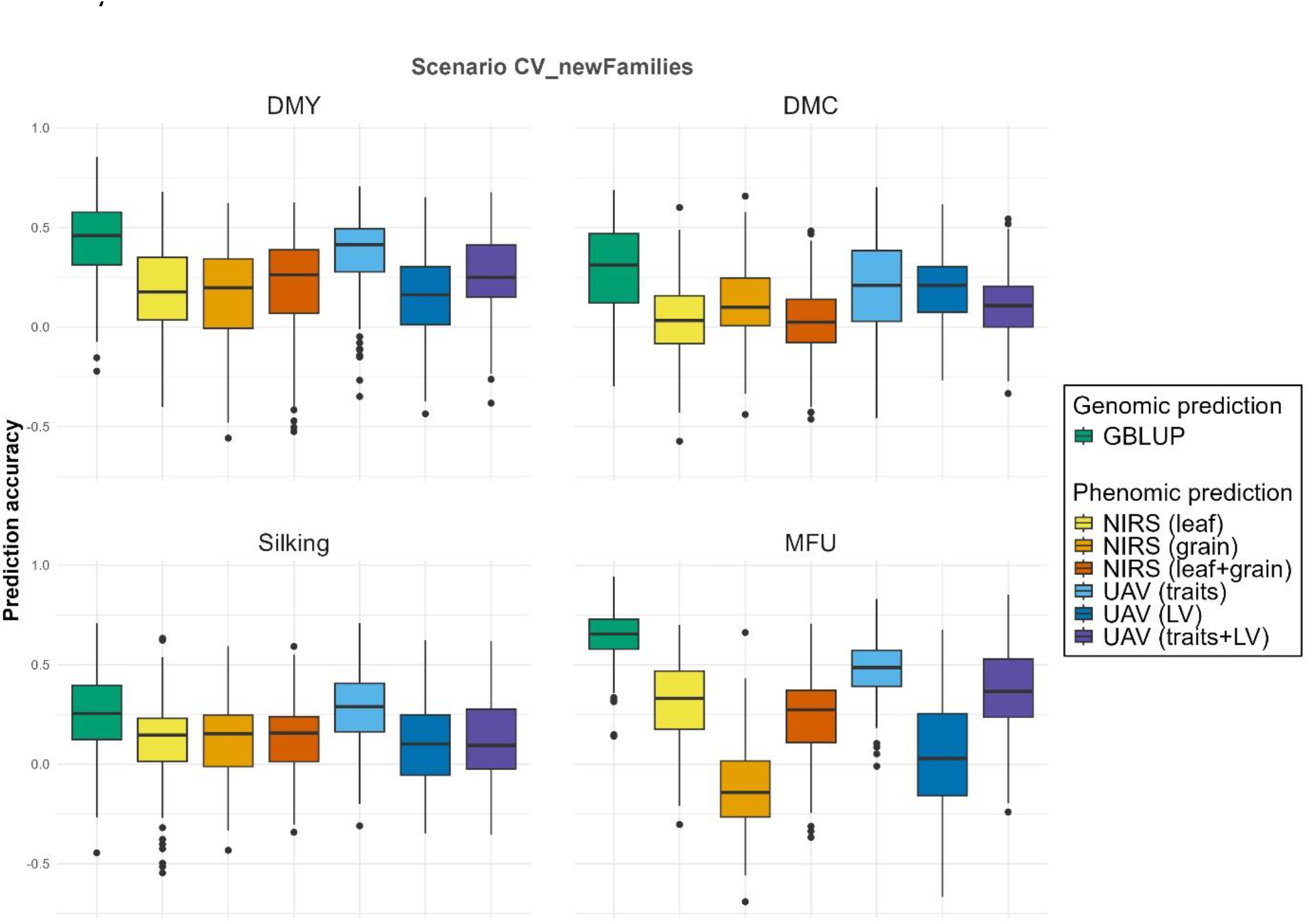
Genomic and phenomic prediction accuracies of genetically distant maize hybrids (scenario CV_newFamilies). The parents of the predicted hybrids originate from families not represented in the training set. Phenomic prediction was based on NIRS data acquired from leaf or grain tissue, and on secondary traits (“traits”) or latent variables (“LV”) extracted from UAV images acquired from the dent and flint parental lines grown in nurseries.

#### UAV-derived secondary traits retain genetic signal with fewer variables

To understand why UAV secondary traits were competitive despite their small number, we quantified the genetically structured variation captured by each predictor class (Table 1). Genomic heritability varied both among predictor classes and between the dent and flint nurseries. Median genomic heritability of UAV secondary traits was 0.35 in dent and 0.14 in flint lines, compared with 0.11 and 0.07 for UAV latent variables. For NIR wavelengths, median genomic heritability was 0.30 (dent) and 0.13 (flint) for leaf spectra, and 0.27 in both groups for grain spectra.

**Table 1:** Summary statistics of 1genomic heritabilities and principal component analysis (PCA) of the different types of predictors: NIR spectra acquired from dried leaves or grains, secondary traits (“traits”) and latent variables (LV) from UAV images. The 198 secondary traits (e.g. biomass traits, vegetation indices) were extracter from UAV images acquired on six dates. The latent variables (LV) were extracted from the UAV images on six dates using V-JEPA.

|  | Dent |  |  |  | Flint |  |  |  |
| --- | --- | --- | --- | --- | --- | --- | --- | --- |
|  | NIRS<br>(leaf) | NIRS<br>(grain) | UAV<br>(traits) | UAV<br>(LV) | NIRS<br>(leaf) | NIRS<br>(grain) | UAV<br>(traits) | UAV<br>(LV) |
| No. of variables | 2151 | 1307 | 198 | 1280 | 2151 | 1307 | 198 | 1280 |
| h<sup>2</sup> (first quartile) | 0.16 | 0.07 | 0.23 | 0.01 | 0.07 | 0.09 | 0.01 | 0.00 |
| h<sup>2</sup> (median) | 0.30 | 0.27 | 0.35 | 0.11 | 0.13 | 0.27 | 0.14 | 0.07 |
| h<sup>2</sup> (third quartile) | 0.44 | 0.41 | 0.48 | 0.25 | 0.20 | 0.40 | 0.35 | 0.19 |
| No. with h<sup>2</sup> > 0.4<sup>a</sup> | 761 | 344 | 73 | 105 | 60 | 322 | 43 | 79 |
| No. PCA axes with h<sup>2</sup> > 0.1<sup>b</sup> | 13 | 16 | 15 | 11 | 9 | 12 | 14 | 16 |

<sup>a</sup> Number of variables with a genomic heritability above 0.4
<sup>b</sup> Number of PCA axes with a genomic heritability above 0.1

The predictor classes differed widely in initial dimensionality, from 2,151 leaf NIR wavelengths to only 198 UAV secondary traits. Yet after principal component analysis, the number of axes with genomic heritability above 0.1 fell within a narrow range of 9–16 across predictor classes and heterotic groups. Despite being the smallest set of predictors, the UAV secondary traits produced 15 such axes in dent and 14 in flint lines. These counts exceed those obtained from the 2,151 leaf NIR wavelengths (13 and 9) and are close to the maximum observed for any predictor class. UAV secondary traits therefore provided the most compact representation of genetic variation, capturing a comparable number of genetically structured independent dimensions from roughly one-tenth as many variables.

#### Three well-spaced flights capture most predictive information

Single-date accuracy depended strongly on acquisition timing (Fig. 4). The first three flights (28 June, 11 July and 24 July) occurred before flowering and generally gave lower accuracies. Accuracy improved after the onset of flowering in the dent nursery, and for most traits the highest single-date accuracies were obtained on 13 August or on 2 October. In CV_newFamilies, late-season flights alone gave accuracies that could be comparable to, and in some cases higher than, those obtained by combining all six dates.

**Figure 4:**
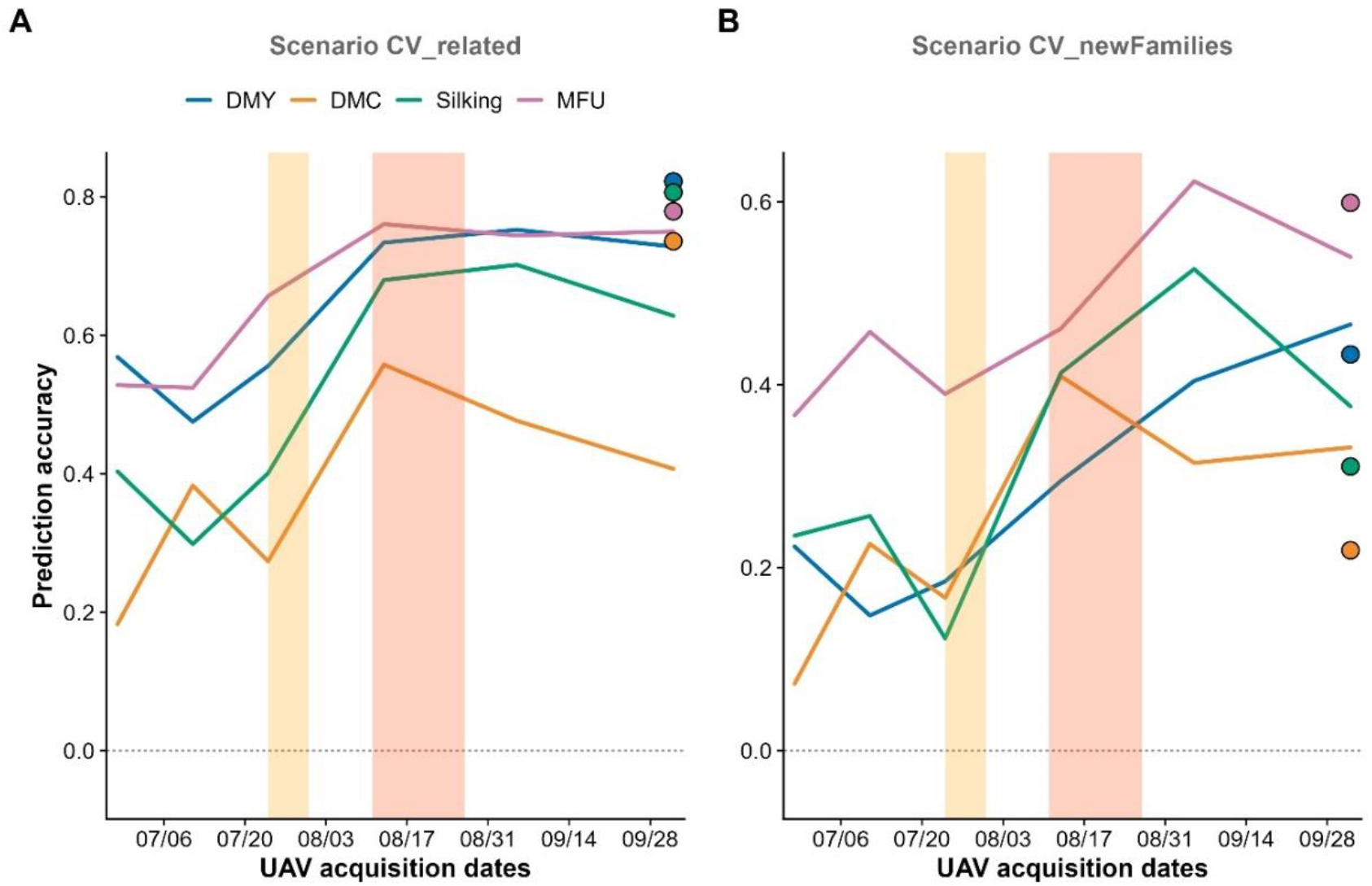
Prediction accuracy of phenomic selection with UAV secondary traits obtained on different dates in maize nurseries. UAV images of parental lines in the nursery were used to predict the performance of the resulting hybrids in field trials. A: results from the random cross-validations (CV_related). B: results obtained when predicting hybrids whose parental lines belong to families not represented among the parents of the training hybrids (CV_newFamilies). The accuracies obtained using the combination of all flights are shown as dots on the right. The yellow and red columns correspond to the silking period of the flint and dent nurseries, respectively.

In CV_related, accuracy increased as acquisition dates were combined, with diminishing gains as more dates were added (Fig. 5). DMC and silking benefited most from additional flights, whereas three dates were sufficient to approach six-date accuracy for DMY and MFU. The gain from combining dates was smaller in CV_newFamilies, where three to four flights approached maximum accuracy for most traits and MFU was only weakly affected by the number of dates (Fig. 6). In both scenarios, a combination of one early, one intermediate and one late flight retained most of the predictive information of the full time series, with accuracies similar to those with four to six flights for CV_related, and to six flights for CV_newFamilies.

**Figure 5:**
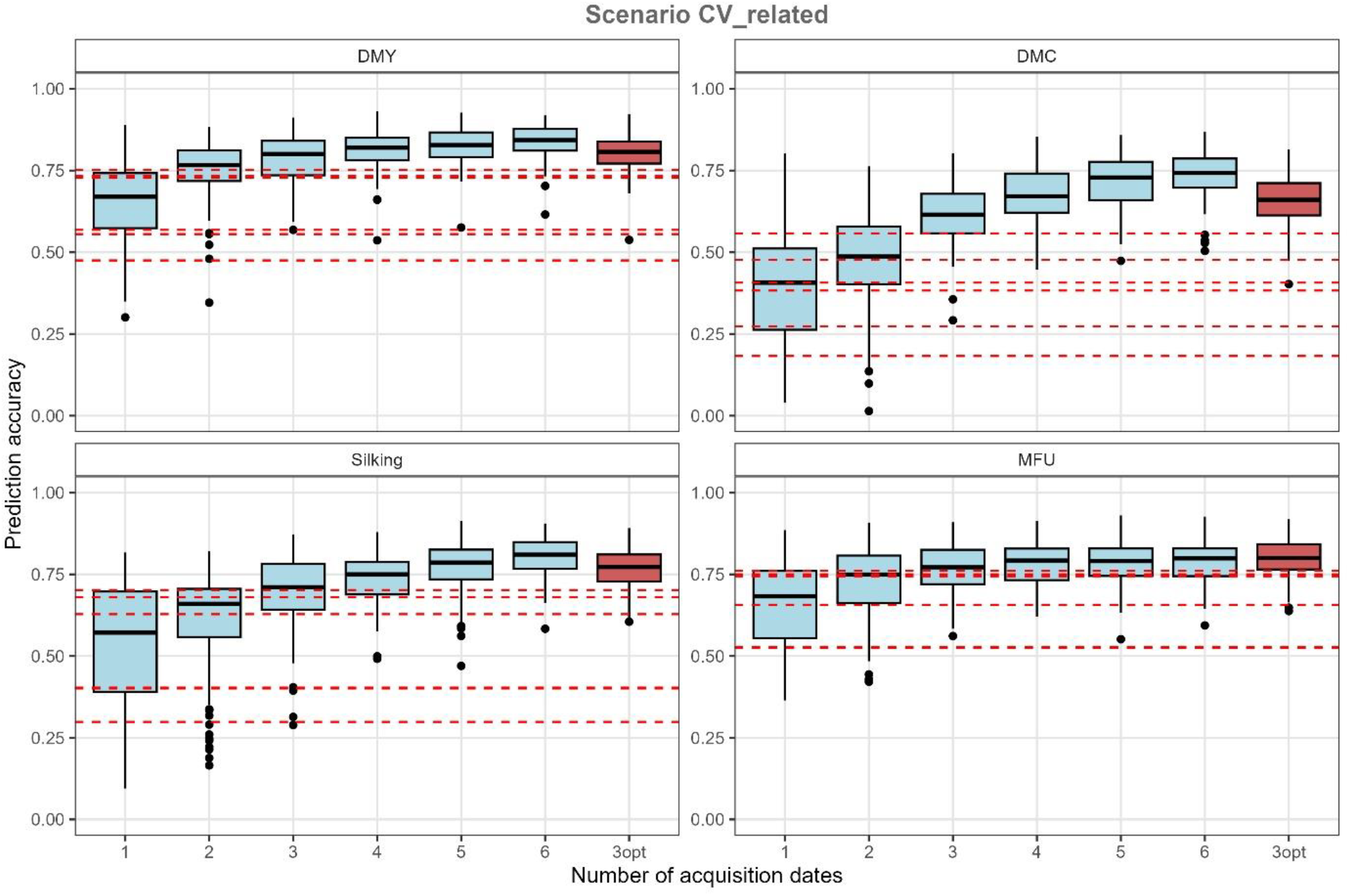
Prediction accuracy of phenomic selection based on UAV images (secondary traits) using combinations of one to six acquisition dates in a random cross-validation scenario (CV_related). 3opt corresponds to the combination of three flights separated in time: an early, an intermediate and a late flight. UAV images of parental lines in the nursery were used to predict the performance of the resulting hybrids evaluated in field trials. The prediction accuracies obtained with each of the six individual flights are shown by red horizontal dashed lines.

**Figure 6:**
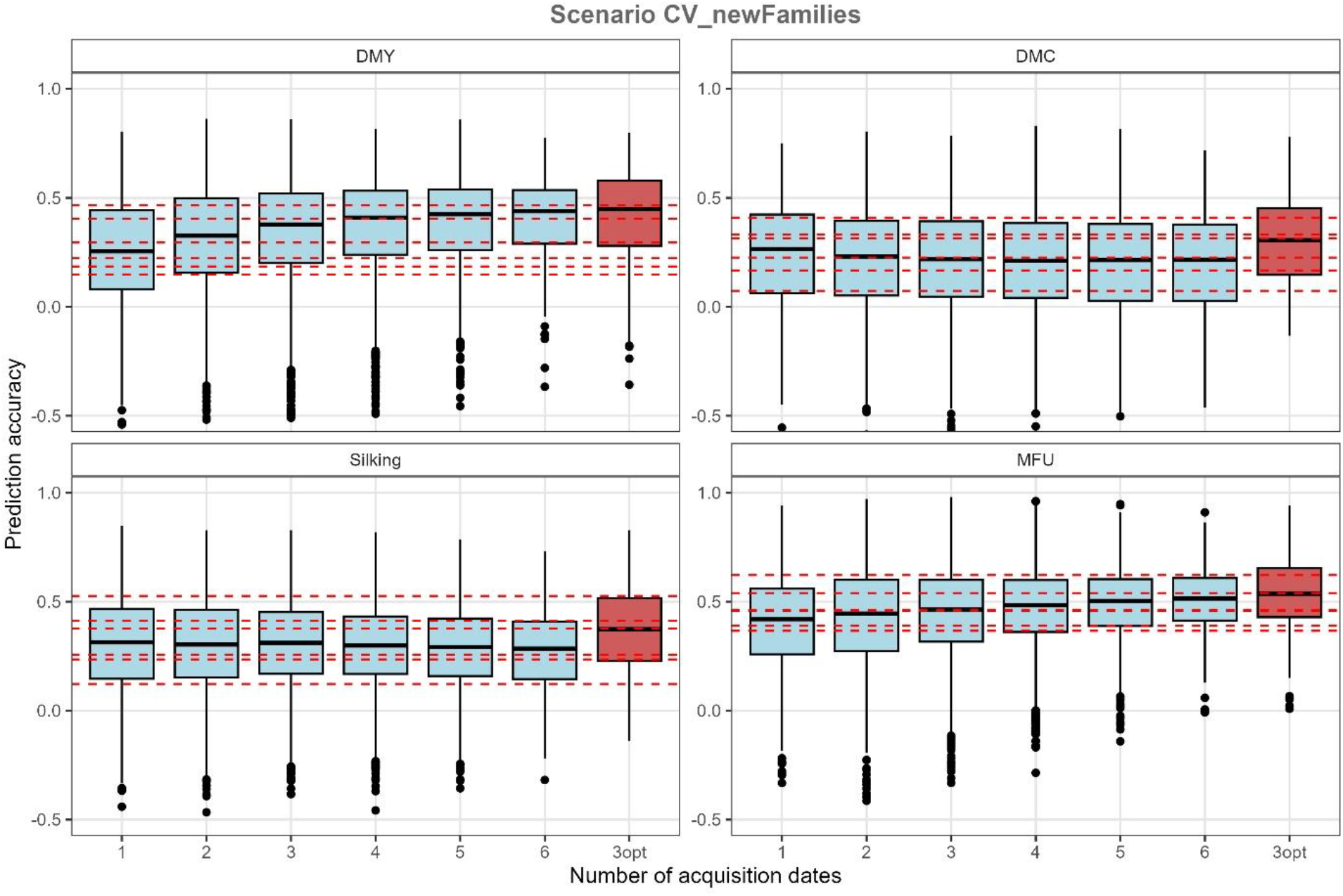
Prediction accuracy of phenomic selection based on UAV images (secondary traits) using combinations of one to six acquisition dates for genetically distant hybrids (CV_newFamilies). 3opt corresponds to the combination of three flights separated in time: an early, an intermediate and a late flight. UAV images of parental lines in the nursery were used to predict the performance of the resulting hybrids evaluated in trials. The prediction accuracies obtained with each of the six individual flights are shown by red horizontal dashed lines.

#### Nursery-based phenomic prediction transfers to winter pea

Pea differs from maize in biology, breeding system and canopy architecture. To assess the generalizability of our approach, we asked whether UAV imagery acquired over a pea nursery could predict the field-trial performance of the imaged genotypes. To this end, we applied the phenomic selection approach to a winter pea nursery comprising 464 genotypes from 100 F4 families. UAV images of the nursery were acquired on four dates, and the phenomic prediction model was trained using an independent field trial. We evaluated the prediction approach using two validation schemes relevant to breeders. In CV_acrossF4, ten F4 families were predicted using the remaining 90 as the training set, with 200 repetitions. In CV_newF4, individual genotypes from an F4 family absent from the training set were predicted. This second scenario is more demanding, because genetic variance within a family is lower than across families. For pea, there was no genomic benchmark because the breeding material had not been genotyped.

In CV_acrossF4, combining UAV secondary traits and latent variables gave accuracies of 0.47 for yield, 0.69 for height and 0.70 for flowering date (Table 2). Secondary traits alone were more predictive than latent variables for yield (0.30 vs 0.15) and height (0.65 vs 0.30), whereas the two representations performed similarly for flowering (0.63 vs 0.65). Combining them increased accuracy for all three traits.

**Table 2:** Phenomic prediction accuracies when predicting the trial performance of genotypes grown in a winter pea nursery. In CV_acrossF4, the training set is composed of 90 F4 families and the test set of the 10 remaining F4 families. In CV_newF4, the test genotypes originate from one F4 family absent from the training set. The predictors were secondary traits from UAV images (“traits”) or latent variables from UAV images (“LV”).

|  | CV_acrossF4 |  |  | CV_newF4 |  |  |
| --- | --- | --- | --- | --- | --- | --- |
|  | Yield | Height | Flowering | Yield | Height | Flowering |
| UAV (traits) | 0.30 | 0.65 | 0.63 | 0.40 | 0.44 | 0.55 |
| UAV (LV) | 0.15 | 0.30 | 0.65 | 0.11 | -0.15 | 0.42 |
| UAV (traits+LV) | 0.47 | 0.69 | 0.70 | 0.37 | 0.43 | 0.56 |

Prediction remained possible within new F4 families (CV_newF4, Table 2). With secondary traits, accuracies were 0.40 for yield, 0.44 for height and 0.55 for flowering. Latent variables were much less predictive (0.11, ™0.15 and 0.42, respectively). Combining both representations gave 0.37, 0.43 and 0.56, providing no consistent improvement over secondary traits alone.

#### Observed accuracies support substantial nursery culling

We then evaluated how much of a nursery can be discarded, at a given prediction accuracy, while controlling the risk of losing superior genotypes. To assess this, we simulated nurseries of 1,500 genotypes (Fig. 7). For each combination elimination rate - prediction accuracy, we computed the risk of having discarded the best genotypes (Fig. 7a), and the proportion of the best 30 genotypes that has been discarded (Fig. 7b).

**Figure 7:**
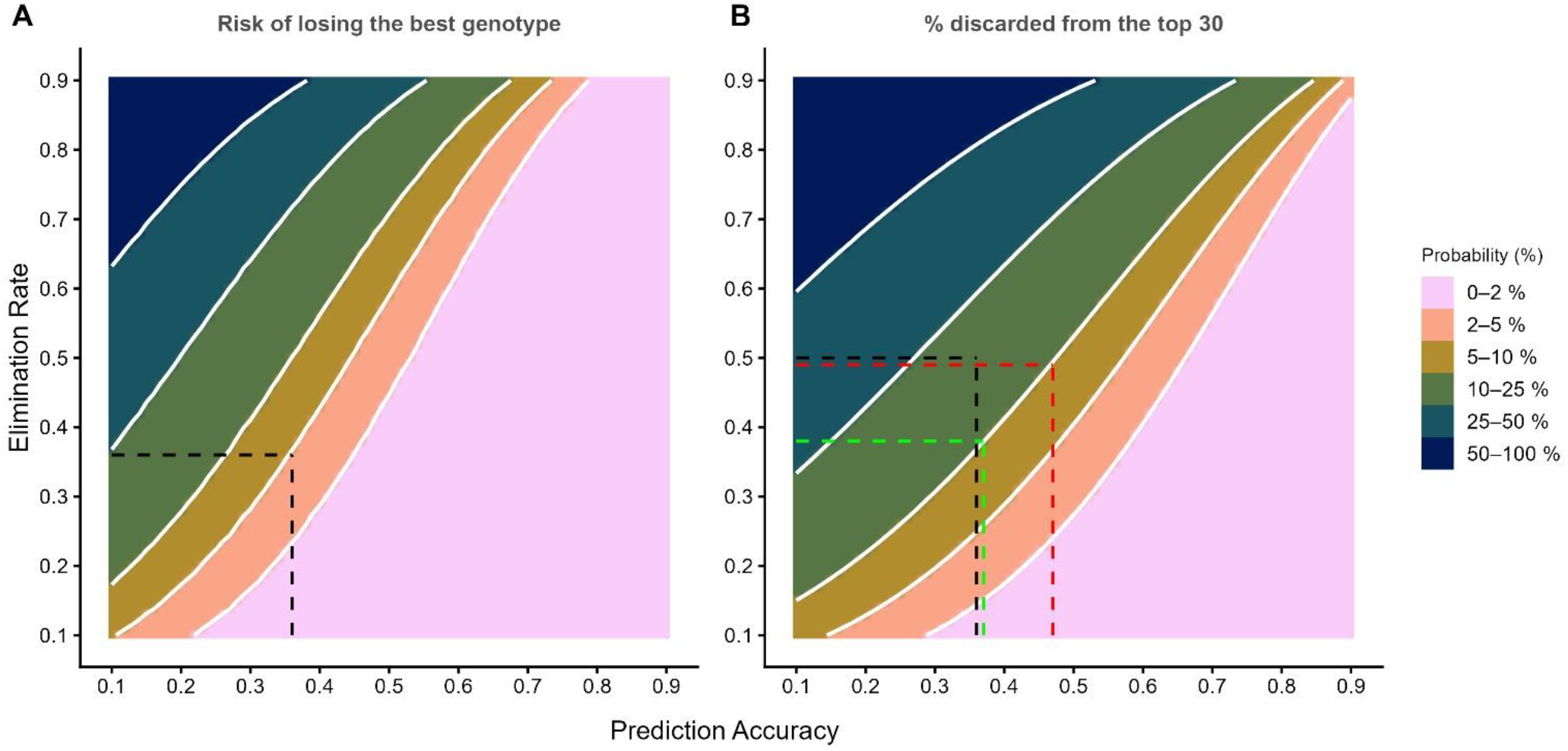
Risk of eliminating the best genotypes for a given elimination rate and prediction accuracy for a nursery of 1,500 genotypes. A: risk of eliminating the best genotype. B: proportion of the top 30 genotypes discarded. Prediction accuracy is the correlation between the predictions and the true genetic values, usually estimated as the predictive ability divided by the square root of the trait heritability. Dashed lines correspond to the maize and pea examples presented in the text (black for maize, green and red for pea)

With the probability of eliminating the single best genotype held below 5%, an accuracy of 0.45 allowed 50% of the nursery to be eliminated (Fig. 7a). At the same risk threshold of 5%, the DMY accuracy of 0.36 observed in maize under CV_newFamilies allowed an elimination rate of 36%. Breeders are often interested in retaining a group of top candidates rather than a single best genotype, so we also quantified losses among the 30 best genotypes (Fig. 7b). At an accuracy of 0.50, 53% of the nursery could be eliminated with an average loss of three of the top 30 genotypes. At the maize DMY accuracy of 0.36, eliminating 50% of the nursery resulted in an average loss of 5.3 of the top 30.

For pea, the yield accuracy obtained in CV_acrossF4 allowed 49% of the nursery to be eliminated with an average loss of three of the top 30 genotypes. The yield accuracy in CV_newF4 allowed 38% to be eliminated at the same average loss. These results should be interpreted cautiously because the simulations did not account for variation in estimated accuracy across cross-validation repetitions.

## Discussion

We show that high-throughput phenotyping data acquired in breeding nurseries can predict the field performance of new selection candidates before they enter field trials. Phenomic selection was initially proposed as a low-cost alternative to genomic prediction based on the observation that high-dimensional phenotypes capture genetic similarities among individuals. Subsequent studies across diverse crops established the general predictive value of this approach. The present work addresses a different question: whether this predictive principle can be translated into a practical selection tool at the nursery stage of breeding programs, where large numbers of candidates must be screened before productivity can be reliably evaluated. We assessed in two complementary species, the ability of UAV imagery acquired over nurseries to predict the field-trial performance of the imaged genotypes. Maize and pea are selected under markedly different breeding systems. Maize is an allogamous species cultivated as hybrids, whereas nurseries consist of parental inbred lines. The objective is therefore to predict the hybrid value of the parental lines grown in nurseries. Pea, by contrast, is an autogamous species, in which both nursery genotypes and field-trial genotypes are inbred lines, resulting in a less challenging phenomic selection scenario than in maize. For both species, the objective is to predict the field performance of genotypes grown exclusively in the nursery.

In maize, UAV imagery acquired on parental inbred lines grown in nurseries predicted the performance of their hybrids in multi-environment trials, including when the parental families of the predicted hybrids were absent from the training population. In winter pea, nursery imagery also predicted yield, height and flowering in a field trial, despite major differences from maize in breeding system, plant architecture and crop management. In the most demanding prediction scenarios, accuracies for yield obtained with UAV secondary traits remained at least 0.36 (0.36 for maize and 0.40 for pea). These accuracies are insufficient to reliably identify the very best genotypes. However, our simulations show that they can support substantial early culling while controlling the risk of discarding superior material. These results establish nursery-based phenomic selection as a potential additional source of information at a stage of breeding programs where selection intensity is high but information on future field performance is limited.

Previous studies have shown that high-dimensional phenotypes can predict complex traits in several crop species^17,19–32,34–37^, including hybrid performance using phenotypes measured on parental lines^26,30^. In these studies, predictions concerned genotypes that had already entered field trials, restricting the scope of phenomic selection to applications such as sparse testing at a stage when many candidates had already been discarded. Extending phenomic selection to nursery screening presents an additional challenge: nursery plants are grown under conditions that differ substantially from those in field trials, including lower planting densities, fewer plants per genotype arranged in rows, and different management practices. The present study provides a proof of concept that phenomic selection can be used as a nursery-screening strategy. Here, the objective is not to predict a trait expressed by the plants being imaged, but to predict their field performance, or that of their hybrid progeny, in independent trials.

In maize, genomic prediction of hybrids from parental genotypes can rely on well-established combining-ability frameworks^42,43^. For phenomic prediction, the parental phenotypes must themselves carry genetic information about hybrid performance. The maize design provides a stringent test of this: UAV and NIR data were collected from parental lines in 2024, whereas hybrid performance was measured across eight trials conducted in 2016 and 2017. In maize, prediction therefore cannot result from environmental effects shared between the nursery and the target field trials. This addresses an important concern in phenomic prediction, because correlations between high-dimensional phenotypes and target traits measured in the same environment can contain substantial non-genetic components^44,45^. The accuracies observed here instead imply that phenotypes expressed in the nursery capture persistent genotype-related differences. These differences are informative about performance in other environments, in a different generation and in a different genetic background. This does not imply that individual UAV traits are causal determinants of hybrid performance, but rather that their joint variation provides a phenomic measure of similarity that is useful for prediction.

The comparison with genomic prediction in maize also helps define the role of this approach. When training and predicted hybrids were closely connected, UAV-based phenomic prediction approached genomic prediction, with mean accuracies of 0.82 and 0.88, respectively. As expected, both approaches lost accuracy when the parental families of the predicted hybrids were excluded from training. Nevertheless, UAV secondary traits retained an accuracy of 0.36 for dry matter yield.

The relevance of these results is not that phenomic selection should replace genomic selection. Molecular markers provide information that phenomic data cannot substitute for, including information for optimizing crosses^46–48^, managing genetic diversity^49–51^ and identifying genomic regions controlling traits. Instead, nursery phenomic selection could reduce the need to genotype every early-generation candidate. UAV-based predictions could be used for an initial, relatively conservative elimination step, with genotyping and more intensive field evaluation then concentrated on the smaller set of candidates that progress through the breeding program. Such a combination could be particularly useful in crops and breeding programs for which recurrent genotyping of thousands of candidates remains a major constraint. This approach could also be useful for major crops, for which the large number of genotypes produced each year entails substantial genotyping costs.

Among the phenomic predictors, relatively simple UAV-derived secondary traits performed particularly well. Despite comprising only 198 variables, they captured a number of genetically structured principal components similar to, or greater than, those captured by the much larger sets of NIR wavelengths and latent image variables. They also consistently outperformed the latent representation in the more demanding prediction scenarios. One possible explanation is that secondary traits were more heritable than latent variables and carried less redundant information than NIR spectra.

NIR spectra were less predictive than UAV imagery in this study, although NIR spectroscopy has provided accurate phenomic predictions in many previous studies^17,19–23^. This difference likely reflects, at least in part, how the spectra were sampled here. Leaf spectra were acquired from one plant per row on a single date, whereas UAV imagery integrated entire rows at several developmental stages. Improved sampling and preprocessing^52^ could therefore raise NIR-based accuracies. Even so, UAV imagery retains a decisive practical advantage at nursery scale, because it requires no tissue sampling and can cover thousands of plots in a single flight.

Likewise, the lower performance of latent variables should not be interpreted as evidence that representation learning is intrinsically less suited to phenomic selection. The video encoder used here was pretrained on generic videos and applied with frozen weights, without adaptation to plant or nursery imagery. Moreover, combining secondary traits and latent variables improved accuracy in some of the less stringent scenarios, indicating that the two representations contain partly complementary information. Larger video encoders^53^, together with plant-specific or nursery-adapted self-supervised pretraining, may therefore extract additional predictive information from image sequences without requiring manual annotation.

The temporal analyses have practical implications for deployment at nursery scale. Three acquisitions distributed across early, intermediate and late developmental stages retained most of the predictive information of the complete six-flight time series, indicating that dense temporal sampling is not required (Fig. 5-6). This matters because the value of UAV-based phenomic selection depends on its ability to scale to nurseries containing thousands of plots^54,55^. Under the acquisition and processing assumptions considered here, three flights would cost less than US$1 per plot for nurseries of 10,000 plots, and less than US$3 per plot for nurseries of 1,000 plots (Supplementary Note 2; see also Reynolds et al.^56^). For a low-risk approach with five flights, the costs would still be below US$5 per plot. These estimates are intended to illustrate the order of magnitude and will vary with nursery size, equipment, labour costs and image-processing workflows. Moreover, UAV acquisition also provides phenotypes related to development, phenology, canopy structure and biomass, that are useful to breeders, so part of these phenomic selection costs can be shared with routine nursery phenotyping.

The organization of breeding programs provides an opportunity to implement this approach without creating a dedicated training population. Advanced genotypes evaluated in field trials are frequently maintained or multiplied in nurseries, and can therefore be imaged with UAVs alongside newly generated candidates. These advanced genotypes provide field observations for model training, while phenomic similarities measured under common nursery conditions connect them to the candidates to be screened. Characterizing the training and prediction sets under the same environmental conditions is an important strength of the approach, because unlike DNA markers, high-dimensional phenotypes are affected by both genotype and environment.

There is considerable scope for improving the approach. Nursery experimental designs could be optimized not only for seed production and visual selection, but also for repeatable phenomic characterization, thereby providing more reliable estimate of phenomic similarity between genotypes.

A substantial body of literature on experimental design optimization could be leveraged for this purpose^57,58^. Approaches that combine high-dimensional phenotypes collected across years and environments from partially overlapping sets of genotypes could further increase the effective size of the training population^22^. Furthermore, many other opportunities exist to improve phenomic selection, by optimizing NIRS data preprocessing^52^, UAV acquisition dates, the number of flights, sensors, image processing, and representation-learning methods.

The simulations clarify how predictions of moderate accuracy could be used operationally. Their primary value is unlikely to be precise ranking of the few candidates ultimately released or used as parents. Instead, they can support negative selection: eliminating candidates with low predicted value, while retaining a sufficiently large fraction of the nursery to limit the probability of losing superior genotypes^59,60^. For example, the yield accuracy in CV_newF4 allowed 38% to be eliminated with an average loss of three of the best 30 genotypes. The appropriate elimination rate will depend on the trait, the breeding stage and the breeder’s tolerance for risk. Prediction accuracy is itself estimated with uncertainty, so operational implementations should use conservative estimates, obtained for example from cross-validation within the training population or using criteria from the mixed model theory such as the prediction-error variance (PEV) or the coefficient of determination^61,62^.

Several limitations need to be addressed before routine implementation. Each species was evaluated in a single nursery season, and prediction was assessed retrospectively rather than through prospective selection of a new breeding cohort. The maize experiment provides strong temporal independence between nursery phenotypes and target field data, but all parental lines were characterized using NIRS and UAV imagery in the same 2024 nursery environment. Future studies should determine how prediction accuracy changes when models are transferred among nursery years, sites and breeding cycles. Prospective experiments, in which candidates are selected using phenomic predictions and subsequently evaluated in field trials, will ultimately be required to quantify realized selection response.

Together, the maize and pea experiments show that phenomic data acquired in a nursery can be used to predict performance beyond the nursery environment. The contrasting biology, breeding systems and canopy structures of the two species suggest that this principle could be transferable to other species. Validation in additional species and breeding cycles will nevertheless be needed to establish its generality. That the approach worked in pea is encouraging in this respect, because pea is a minor crop in Europe with genotyping resources more limited than those of the major cereals. UAV-based phenomic selection therefore provides a scalable complement to existing breeding tools. It does not require genotyping the selection candidates, can be integrated into nurseries already used for seed multiplication and selection, and can predict productivity before candidates enter expensive field trials. Its main opportunity may lie not in replacing genomic selection, but in extending predictive selection to the many species and breeding programs for which recurrent genotyping of all selection candidates is not currently feasible.

## Acknowledgments

We thank the ProMaïs consortium of the SAMMCR project (Lidea, Limagrain Europe, Maïsadour Semences, Corteva, RAGT 2n, and Syngenta Seeds) for inbred lines development, genotyping, and for the phenotyping of the maize hybrid dataset.

## Author contributions

RR, LM and CB designed the experiments. LN, CB, FB, EG and ALM managed the field experiments and nurseries. GT, FB, DG, ALM and EG coordinated the pea trials and nurseries. CB, LM, LN and RR coordinated the maize trials and nurseries. ME supervised NIR spectroscopy, and ME and LN acquired the NIR spectra. PD and AC processed the UAV images, derived the secondary traits and latent variables. RR, QH and AGD performed the statistical modelling and analyses. RR wrote the manuscript. All authors edited and approved the manuscript.

## Funding

The maize study was funded by INRAE and SPS—Sciences des Plantes de Saclay. UAV and NIRS data acquisition for the pea study was supported by the “Compte d’Affectation Spéciale pour le Développement Agricole et Rural” of the Ministère de l’Agriculture, de l’Agroalimentaire et de la Souveraineté Alimentaire de la République Française.

## Competing interests

AC, DG and PD are employees and/or shareholders at HiPhen.

## Data availability

Data are available at https://github.com/rrincent/PhenomicNurseryScreening_Pea for pea and at https://github.com/rrincent/PhenomicNurseryScreening_Maize for maize.

## Code availability

Code is available at https://github.com/rrincent/PhenomicNurseryScreening_Pea for pea and at https://github.com/rrincent/PhenomicNurseryScreening_Maize for maize.

## Methods

### Maize genetic material – Incomplete factorial design

Originally, six connected biparental families were developed within each heterotic group—dent and flint—yielding a total of 12 families ^26,63–65^. Within each group, the six families originated from crosses among four founder lines selected for their combining ability with the other group for silage-related traits, including digestibility and yield. In the flint group, lines were advanced through five to six generations of selfing using a single-seed descent scheme due to lower responsiveness to doubled haploid production, whereas dent lines were derived via doubled haploid production.

An incomplete factorial mating design was subsequently implemented using a subset of 90 flint and 90 dent lines. Under this sparse crossing scheme, each parental line contributed to approximately four hybrids (see Lorenzi et al.^42^ for further details). Overall, this design generated 363 dent × flint hybrids.

### Phenotyping

The hybrids were evaluated across eight field trials conducted over two years, with three trials in 2016 and five in 2017^42^. These experiments were carried out by seven private breeding companies (RAGT, Lidea, KWS, Corteva, Masseeds, Syngenta, and Limagrain) members of the Promaïs SAMMCR consortium. Trials followed partially replicated augmented designs as described by Williams et al. (2011)^66^. Two types of control hybrids were included: two commercial checks (LG30.275 and RONALDINIO) and 16 founder hybrids obtained by crossing the four founder lines within each heterotic group. In each trial, all checks, along with approximately 20% of the experimental hybrids, were evaluated in duplicate.

Hybrids were phenotyped for four agronomic traits—dry matter yield (DMY), dry matter content (DMC), female flowering time (silking), and plant height (PH)—as well as seven silage quality traits: milk fodder unit (MFU), neutral detergent fiber (NDF), digestibility of non-starch and non-soluble carbohydrates (DINAG), digestibility of non-starch, non-soluble carbohydrates, and non-crude protein (DINAGZ), lignin content (LIGN), cellulose content (CELL), and hemicellulose content (HCELL). Silage quality traits were predicted using near-infrared spectroscopy (NIRS) measured at harvest. Consistent with Lorenzi et al.^42^ (2022), subsequent analyses focused on four key traits representing productivity (DMY, DMC), phenology (silking), and quality (MFU).

Details on data curation, estimation of broad-sense heritability, and calculation of adjusted means at the multi-environment trial level are provided in Lorenzi et al.^42^. Broad-sense heritability estimates were high, reaching 0.89 for DMY, 0.93 for both DMC and silking, and 0.86 for MFU.

### Genotyping

All founder and parental lines were genotyped using an Affymetrix® array (provided by Limagrain), yielding 18,480 SNP markers. Marker quality control was performed separately within the dent and flint parental groups. Markers were excluded if they exhibited more than 20% missing data, heterozygosity rates exceeding 5% in dent lines or 10% in flint lines, or a minor allele frequency below 5%. After filtering, a total of 9,548 polymorphic SNPs were retained for downstream analyses. Genotypes were encoded as 0, 0.5, or 1, corresponding to the dosage of an arbitrarily defined reference allele. Missing genotypes were subsequently imputed using AlphaPlantImpute ^67^.

### High-throughput phenotyping of the nursery

Parental lines included in the incomplete factorial design were grown in two nurseries corresponding to the dent and flint heterotic groups, at Gif-sur-Yvette, France, in 2024. The flint nursery comprised 195 rows representing 83 genotypes, with an average of 2.3 rows per genotype, while the dent nursery included 191 rows corresponding to 69 genotypes, with an average of 2.8 rows per genotype. Each row contained approximately 25 plants, although plant density varied substantially due to heterogeneous germination rates among genotypes. The two nurseries were separated by a single row in which the eight founder lines were sown. After applying the quality-control procedures described below, complete data were available for the parental lines of 242 dent × flint hybrids (77 flint parents and 68 dent parents).

For NIRS acquisition on leaves, the last ligulated leaf of one plant per row was sampled on a single date around flowering time. Leaves were immediately pressed to maintain flatness and subsequently oven-dried at 50 °C for 72 hours. Near-infrared (NIR) spectra were acquired from dried leaves using a FieldSpec 2500® portable spectrometer (Analytical Spectral Devices, Inc., Boulder, CO, USA) equipped with a bifurcated fiber-optic probe. For each sample, spectra were recorded at two positions on the adaxial surface, on either side of the midrib. Reflectance calibration was performed using a Spectralon® reference panel, which was also placed beneath the leaf during acquisition. The spectrometer covered a wavelength range of 350–2500 nm, with a spectral resolution of 3 nm in the 350–1000 nm region and 10 nm in the 1000–2500 nm region. The sampling intervals were 1.4 nm and 2 nm for these respective ranges. During measurements, leaves were positioned with the adaxial surface in direct contact with the light source, and the fiber-optic probe was connected to the detector. Harvested grains were dried at 40 °C for 72 hours. NIR spectra were acquired from the dried grains using a Perkin Elmer FT9700, with a spectral resolution of 16 cm^-1^ over the 3856-14,304 cm^-1^ range, averaging 32 scans per sample.

Aerial images were acquired on six dates spanning the full maize growing season, from plant emergence to senescence (28 June, 11 July, 24 July, 13 August, 5 September and 2 October 2024). On each date, two successive flights were conducted over the nursery under clear-sky conditions between 11:00 and 13:00 local time, when the solar elevation angle exceeded 30°, to minimize shadow effects. A detailed description of the acquisition procedure is provided in the Supplementary Note 3.

### Processing of NIR spectra and UAV-derived data

NIR spectra were first normalized by centering and scaling. For each wavelength, spatial variation was accounted for by fitting a smoothing spline model using the R package *SpATS*^68^, with row and column coordinates included as random effects. Genotype was modeled either as a fixed effect to estimate adjusted marginal means or as a random effect to derive broad-sense heritability.

The resulting adjusted means were subsequently re-standardized, and their first derivative was computed using the Savitzky–Golay filter^69^, with a window size of 37 data points and a polynomial order of 2, as implemented in the R package *signal* (Signal Developers 2014).

### Processing of UAV images (secondary traits)

For each plot, a maximum of 44 secondary traits were extracted on each date from raw RGB and multispectral images after radiometric calibration, using the Hiphen pipeline. Traits were organized into six categories: canopy development (e.g., VARI), trial quality and harvest index (e.g., plant count and density), biophysical variables (e.g., FiPAR), multispectral vegetation indices (e.g., NDVI and NDRE), raw reflectance values in the four multispectral bands, and biomass proxies (e.g., plant height and biovolume). A detailed description of these secondary traits is provided in the Supplementary Note 4. As for the spectra, a smoothing-spline spatial model was fitted to each secondary trait.

### Processing of UAV images (latent variables)

Individual plants were first detected within each UAV image and spatially grouped into triplets of three consecutive plants. Each triplet was cropped using a bounding box held constant across all six acquisition dates, ensuring that the same three plants were followed throughout the season. The six crops of a given triplet were resized to 384 × 384 pixels and stacked along the temporal dimension, forming a 6 × 384 × 384 array that was treated as a video clip.

Embeddings were extracted using V-JEPA^38^, a self-supervised video model based on a joint-embedding predictive architecture. We used the publicly available V-JEPA weights (ViT-H/16 backbone) without any fine-tuning, taking the checkpoint released for the input resolution used, and ran the encoder in inference mode with frozen weights. Clips were tokenized with a spatial patch size of 16 × 16 pixels and a temporal tubelet size of 2 frames, yielding 1,728 spatio-temporal tokens of dimension 1,280 for the maize clips. The tokens were then pooled into a single 1,280-dimensional embedding vector per triplet by an untrained attention-based pooling layer, with weights initialized once using a fixed random seed and shared across. A smoothing-spline spatial model (SpATS) was fitted to each latent variable.

### Computation of the genomic and phenomic covariance matrices

Genomic kinship matrices for the dent and flint general combining ability (GCA, 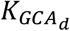and 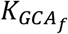) were computed from SNP genotypes for all parental lines using the first method described by VanRaden ^70^, adapted for inbred lines. The kinship coefficient of the dent GCA between individuals i and i’ was estimated as:

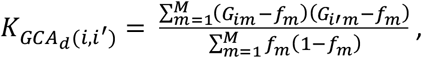

where *G*_*im*_ denotes the genotype of dent line i at locus m (coded 0, 0.5 and 1 for BB, BA and AA genotypes) and *f*_*m*_ denotes the allele frequency of the reference allele A at locus m estimated on the dent parental lines. The kinship matrix 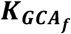 for the flint parental lines was computed similarly.

For NIR spectra, UAV secondary traits and latent variables, the covariance matrices, also referred to as H matrices in the literature, were computed as described in Robert et al. ^23^:

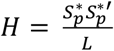, where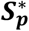denotes the centered and scaled matrix of preprocessed variables, and L is the number of variables. The H matrix was then split into two matrices, one for the dent lines 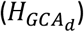 and one for the flint lines 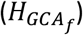.

All genomic and phenomic covariance matrices were scaled by dividing them by the average of the diagonal elements.

### Genomic and phenomic prediction models

Different prediction models relying on genomic or phenomic covariance matrices were defined to compare the accuracies of the different predictors and their potential complementarity. The models were as follows:

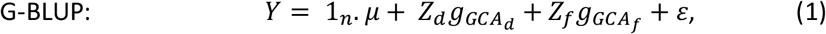

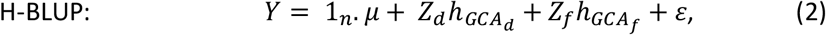

where ***Y*** is the vector of adjusted phenotypic means across the eight hybrid trials, *μ* is the intercept, and 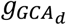 is the random genomic GCA-effect vector for the dent (flint) parents, with 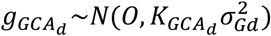 and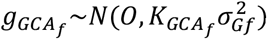. ***Z***_***d***_ and ***Z***_***f***_ are the corresponding design matrices. *ε* is the residual-error vector, with 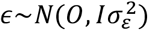, and I is the identity matrix. No specific combining ability (SCA) effect was included in the models as it did not improve predictions.

Similarly, 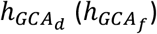 is the random phenomic GCA-effect vector for the dent (flint) parents, with 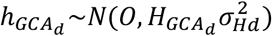and 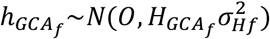, the covariance matrices 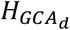 and 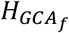 being estimated with phenomic data (NIR spectra, secondary traits from UAV images, or latent variables), as described above.

Different phenomic effects were also combined into a single model:

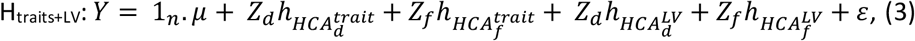

with 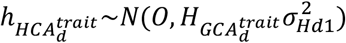 and 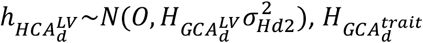being the covariance matrices among the dent parents estimated with the secondary traits and the latent variables extracted from the UAV images, respectively.

### Prediction scenarios and the corresponding cross-validation schemes

We evaluated the predictive performance of genomic and phenomic models across two scenarios.

In CV_related, the 242 hybrids were randomly partitioned into a training set (80%) and a predicted set (20%), without accounting for population structure among parental lines. This process was repeated 200 times.

In CV_newFamilies, the objective was to evaluate prediction accuracy in a situation in which the training set and the predicted set are genetically distant. For this, we accounted for the origins of the dent and flint parental lines from six families each. Training and predicted sets were constructed such that the parents of the predicted hybrids belonged to families distinct from those represented in the training set, with the constraints that at least 20 hybrids were in the validation set and at least 120 hybrids were in the training set, leading to 150 partitions. The UAV secondary traits and latent variables with a broad-sense heritability (computed without genotyping data) below 0.4 were removed.

In these two scenarios, we compared the effect of the acquisition dates and of the number of acquisition dates on prediction accuracy. We also tested the interest of combining an early (11 July), an intermediate (13 August) and a late date (2 October). These three dates were chosen based on their proximity with key stages (early growth, flowering and maturity). All models were evaluated on identical partitions. Predictive performance was measured for each fold as the Pearson correlation between predicted values and adjusted means in the prediction set divided by the square root of the trait heritability.

### Analysis and comparison of the phenomic predictors

To estimate genomic heritability, a simple genomic best linear unbiased prediction model was applied to each NIR wavelength, each secondary trait, and each latent variable. For each type of predictor (wavelengths, secondary traits, latent variables), the number of predictors with a genomic heritability above 0.4 was computed.

A principal component analysis (PCA) was applied to each type of predictor, and we computed the number of axes with a genomic heritability above 0.1 to quantify the amount of genetic information.

### Validation in a pea nursery

To validate the approach in pea, we used the 2025 winter nursery of the Florimond-Desprez breeding program, and the corresponding field trial. UAV images were acquired on four dates (13 March, 30 April, 23 May and 14 June 2025). The nursery comprised 464 genotypes from 100 F4 families, with two adjacent rows per genotype. These genotypes were also evaluated in an independent field trial but the same year, with two replicates per genotype. The UAV platforms and flight protocol followed those used for the maize nurseries. Two aspects of the image processing differed. First, the unit of encoding was the microplot rather than a group of plants: a single video clip was built per microplot, with one frame per acquisition date obtained as a square crop centred on the microplot and resized to 224 × 224 pixels. Second, the V-JEPA checkpoint released for that resolution was used. The temporal tubelet spans two frames, so sequences with an odd number of acquisition dates were padded to an even length: the four dates yielded four frames, corresponding to 392 tokens per clip respectively. Tokens were pooled into a single 1,280-dimensional embedding vector per microplot using the same untrained pooling layer as for maize. The resulting secondary traits and latent variables, as well as the trial phenotypes (yield, plant height before harvest, and onset of flowering) were corrected for spatial effects using SpATS^68^. More precisely, the genotypes originating from the same F4 family were grouped to improve the correction for spatial effects. The phenomic prediction model was the same as for maize, with a phenomic similarity matrix estimated with the secondary traits or the latent variables of the four acquisition dates. We estimated prediction accuracy in two complementary scenarios relevant to breeders: the prediction of 10 new F4 families using the remaining 90 F4 families (CV_acrossF4), and the prediction of genotypes derived from a single F4 family not represented in the training set (CV_newF4). For CV_acrossF4, the partition process was repeated 200 times, and for CV_newF4 we predicted only the F4 families with at least ten genotypes (11 families). For CV_newF4, because we needed to predict the variability within F4 families, spatial correction of the nursery UAV traits was performed at the genotype level (rather than at the F4-family level, as in the CV_acrossF4 scenario).

The prediction accuracies were computed as the Pearson correlation between the predictions and the adjusted phenotypes divided by the square root of the trait heritability.

### Evaluation of risk when screening nurseries with predictive approaches

For a given accuracy, we simulated nurseries of 1,500 genotypes and generated true genetic values and the corresponding predictions using a bivariate normal model with a correlation between the two variables equal to the accuracy. Selection was then applied to predictions for a given elimination rate. This simple simulation framework was repeated 500,000 times. For each accuracy - elimination-rate combination, we then computed, across the 500,000 repetitions, the average probability of losing the best genotype, and the average proportion of discarded genotypes among the top 30. The results were presented in the form of risk maps.

## Notes

https://github.com/rrincent/PhenomicNurseryScreening_Pea

https://github.com/rrincent/PhenomicNurseryScreening_Maize

