## Supplementary Notes for "Phenomic selection enables nursery screening for field performance in maize and pea"

### Supplementary information

#### Supplementary Note 1: Agronomic traits heritabilities

Broad sense heritabilities for the maize hybrid panel were of 0.89 (DMY), 0.93 (DMC), 0.93 (Silking), and 0.86 (MFU). For pea, they were of 0.81 (Yield), 0.68 (plant height) and flowering (0.96).

#### Supplementary Note 2: Cost of UAV images acquisition and treatment

1. Economic analysis by Reynolds et al. (2019) [11]

Based on Table 2, we estimate that the total cost of 10 UAV acquisition dates for 1,700 plots is $12,400. These costs include image capture, image analysis, and data storage for ten years. We excluded agronomic costs ("microplots or plant handling"), because the nursery would be sown and managed regardless. We also excluded trait-analysis costs, because the image-derived outputs are used directly for phenomic prediction. The resulting cost is $0.73 per plot per flight, corresponding to $2.20 per plot for three acquisition dates and $3.65 per plot for five acquisition dates.

2. HiPhen internal costs

The cost structure comprises three components for each acquisition date and nursery: depreciation of the UAV platform (a consumer-grade multispectral system such as the DJI Mavic 3M, €6,000–8,000, shared across sites and depreciated over three years, i.e. €50–90 per acquisition date), one technician-day for field operations (€200–600 depending on the country), and image processing at commercial service rates. The latter term is the most variable because it depends on both nursery size and overall contract volume: it ranges from €500 to €1,500 per date for small, one-off projects, and from €300 to €750 per date for large multi-site, multi-date contracts, for which processing costs are substantially lower per unit.

For a nursery of 10,000 genotypes screened at three acquisition dates under a large-contract regime, these costs correspond to approximately €0.2–0.4 per genotype. For a small, one-off project involving a nursery of 1,000 genotypes, the cost is approximately €3 per genotype. Adopting a lower-risk strategy with five acquisition dates results in costs of approximately €0.4–0.6 and €5 per genotype, respectively. The reduction in per-genotype cost with increasing nursery size therefore arises from two compounding effects: the fixed per-date costs are spread over ten times more candidates, and the per-date processing cost itself decreases with contract volume. These figures correspond to routine implementation in nurseries with usable plot metadata; costs would be substantially higher in a proof-of-concept study such as the present one, with fewer genotypes and non-optimised pipelines.

We do not account for the fact that image-derived traits are of interest to breeders in their own right (e.g., height, ground cover, early vigour, and biovolume), meaning that the full cost is not solely attributable to phenomic selection.

#### Supplementary Note 3: Description of the UAV image acquisition procedure

For maize, high-resolution RGB imagery was acquired using a DJI Matrice 300 RTK UAV (DJI, Shenzhen, China) equipped with a Zenmuse P1 camera (35 mm focal length, full-frame sensor), flying at 20 m above ground level and yielding a ground sampling distance (GSD) of 2.5 mm pixel$^{-1}$. Multispectral imagery was acquired on the same dates using a DJI Mavic 3 Multispectral UAV equipped with a 20-megapixel RGB camera and four co-registered 5-megapixel single-band cameras covering green (G,

550 ± 16 nm), red (R, 650 ± 16 nm), red-edge (RE, 730 ± 16 nm) and near-infrared (NIR, 860 ± 26 nm), flying at 13 m above ground level and yielding a GSD of 6 mm pixel$^{-1}$. Both UAVs flew in automated grid patterns with 80% forward and 75% side overlap, in aperture-priority mode to maintain consistent exposure. Before each flight, radiometric calibration of the multispectral camera was performed using a HiPhen calibrated grey reference panel. A bare-soil reference zone - a vegetation-free area declared within the trial layout - was identified at each flight date to provide a ground-level radiometric reference for multispectral correction, following HiPhen standard processing pipeline. Five ground control points (GCPs) were deployed across the nursery and georeferenced using an Emlid Reach RS2+ GNSS receiver (Emlid, Hong Kong) to ensure consistent spatial co-registration across all flight dates. Images were processed using Agisoft Metashape Professional (generated by Phenoscript 1.11.1, Agisoft LLC, St. Petersburg, Russia) to generate co-registered orthomosaics and dense point clouds via structure-from-motion (SfM) photogrammetry. Plot-level trait values were extracted directly from the raw images within georeferenced row boundaries.

For pea, RGB and multispectral imagery were acquired simultaneously with a single DJI Mavic 3 Multispectral (M3M) platform, which carries a 20-megapixel RGB camera and four co-registered 5-megapixel single-band cameras covering green (550 ± 16 nm), red (650 ± 16 nm), red-edge (730 ± 16 nm) and near-infrared (860 ± 26 nm). All flights were performed at 13 m above ground level, giving a ground sampling distance of 3.5 mm pixel$^{-1}$ for the RGB camera and 6 mm pixel$^{-1}$ for the multispectral bands. The platform followed automated grid patterns with 80% forward and 75% side overlap, in aperture-priority mode to maintain consistent exposure, under clear-sky conditions between 11:00 and 13:00 local time with a solar elevation angle above 30° to minimise shadow effects.

#### Supplementary Note 4: Description of the secondary traits extracted from the UAV images

(i) Canopy development: fractional green cover, its area and spatial heterogeneity, derived from RGB image segmentation using a fixed threshold applied to the VARI index[1]; and three RGB-based greenness indices (excess-green, green-to-red ratio and VARI[1,2]), with within-plot heterogeneity quantified for each.

| Index | Full name | Formula |
|---|---|---|
| **ExG** | Excess Green Index | ExG = 2G − R − B |
| **G/R** | Green-to-Red ratio | G/R = G / R |
| **VARI** | Visible Atmospherically Resistant Index | VARI = (G − R) / (G + R − B) |

(ii) Trial quality and harvest index: plant count and density (BBCH 10–30) and panicle count and density (BBCH 60 to maturity), estimated from RGB images using a deep-learning-based Faster R-CNN object-detection algorithm[3–5], together with measures of spatial heterogeneity; plant gap metrics (count, distance, percentage); and row-length metrics (maximum, mean, minimum and sum).

(iii) Biophysical variables: leaf area index, fraction of intercepted PAR (FiPAR), leaf and canopy chlorophyll content, and average leaf angle, estimated by inversion of the coupled PROSPECT–SAIL radiative-transfer model[6,7] applied to multispectral reflectance[8].

(iv) Multispectral vegetation indices: NDVI, NDRE, CI*green*, CI*red-edge*, MTCI, LCI, MCARI and Simple Ratio[9].

| Index | Full name | Formula |
|---|---|---|
| **NDVI** | Normalized Difference Vegetation Index | NDVI = (NIR − R) / (NIR + R) |

| | | |
|---|---|---|
| **NDRE** | Normalized Difference Red-Edge Index | NDRE = (NIR − RE) / (NIR + RE) |
| **CI*green*** | Green Chlorophyll Index | CI*green* = (NIR / G) − 1 |
| **CI*red-edge*** | Red-Edge Chlorophyll Index | CI*red-edge* = (NIR / RE) − 1 |
| **MTCI** | MERIS Terrestrial Chlorophyll Index | MTCI = (NIR − RE) / (RE − R) |
| **LCI** | Leaf Chlorophyll Index | LCI = (NIR − RE) / (NIR + R) |
| **MCARI** | Modified Chlorophyll Absorption Reflectance Index | MCARI = [(RE − R) − 0.2(RE − G)] × (RE / R) |
| **SR** | Simple Ratio | SR = NIR / R |

(v) Raw reflectance: the four multispectral band values (G, R, RE, NIR).

(vi) Biomass proxies — height and biovolume: metrics were derived from the dense three-dimensional point cloud generated by structure-from-motion photogrammetry of the RGB images. Maximum plant height per plot was defined as the 99$^{th}$ percentile of point heights within the plot point cloud[10] mean plant height was defined as the average height of the reconstructed canopy surface above the bare-soil reference zones. Height heterogeneity was quantified using the coefficient of variation (CV) of maximum and mean heights calculated over 30-cm sections along the plot. Biovolume was calculated using a regular grid at the point-cloud resolution (0.025 m cell size) as the sum of the maximum height in each grid cell multiplied by the cell area. These six traits were available only from BBCH stage 30 onward, once sufficient canopy development allowed reliable three-dimensional reconstruction.
